# Adaptive trade-offs between niche-driven defence system selection and horizontal gene transfer suggests clinical success in *Acinetobacter* spp

**DOI:** 10.1101/2025.08.01.668115

**Authors:** Vigneshwaran Muthuraman, Proyash Roy, Paul Dean, Bruno Silvester Lopes, Saadlee Shehreen

**Affiliations:** Science Department, School of Health and Life Sciences, Teesside University, Middlesbrough, TS1 3BX, UK; National Horizons Centre, Teesside University, Darlington DL1 1HG, UK

**Keywords:** *Acinetobacter baumannii*, bacterial defence systems, antimicrobial resistance, horizontal gene transfer, phage therapy

## Abstract

2.

**Introduction:** Bacteria face relentless selective pressure from both antibiotics and bacteriophages, driving the evolution of diverse defence systems and shaping horizontal gene transfer (HGT) dynamics.

**Gap Statement:** The relationship between bacterial defence system repertoires and horizontal gene transfer in clinically relevant *Acinetobacter* species remains poorly understood, limiting our ability to predict resistance evolution and design targeted phage therapies.

**Aim:** To characterize defence arsenal across *Acinetobacter* species and investigate their associations with different HGT markers

**Methodology:** We performed a comparative genomic analysis of 132 genomes from 18 Acinetobacter species, focusing on the interplay between defence architectures and HGT markers.

**Results:** Our result reveals pronounced differences in defence system repertoires between lineages: while most Acinetobacter spp. harbour a multilayered arsenal, the clinically dominant A. baumannii international clone 2 (IC2) exhibits a depleted but specialised defence landscape, notably enriched in the phosphorothioation-based SspBCDE system and nearly devoid of restriction-modification systems. Strikingly, many defence systems display mutual exclusivity, and defence system is tightly linked to the presence of mobile elements, antibiotics, and heavy metal resistance. Plasmid-borne defence systems, especially BREX, are prevalent, highlighting the role of mobile elements in distributing both immunity and resistance traits. These patterns suggest that successful clinical lineages finely balance immune defences and genetic plasticity, facilitating rapid adaptation under antimicrobial selection and phage threats.

**Conclusion:** These findings provide insights into evolutionary trade-offs underpinning multidrug resistance and have implications for designing targeted phage therapies against recalcitrant Acinetobacter infections.

**Data summary:** All bioinformatics analysis workflows, custom scripts, and R code are publicly available at https://github.com/vikos77/Acinetobacter-defence-systems. The repository includes bash scripts for genome downloading and tool execution (scripts 1-10), R scripts for statistical analysis and visualization (scripts 1-9), and conda environment files for reproducible software installation. Complete analysis protocols are documented in the repository README and individual script headers. All software versions and parameters are specified in the code documentation.

**Impact Statement:** This study reveals how defence system composition in Acinetobacter species is shaped by ecological niche and linked to horizontal gene transfer, antibiotic resistance, and mobile genetic elements. Notably, the clinically dominant *A. baumannii* IC2 lineage exhibits a streamlined defence repertoire favouring genetic plasticity, facilitating rapid adaptation under antibiotic and phage pressure. These findings enhance our understanding of resistance evolution and support the rational design of phage-based therapies targeting multidrug-resistant pathogens.

## Introduction

Bacteria are frequently challenged by bacteriophages (viruses infecting bacteria) and mobile genetic elements (MGEs), driving the evolution of a wide range of defence systems. These systems are often clustered in specific genomic regions known as “defence islands”. They include well-known mechanisms such as CRISPR-Cas and restriction-modification (R-M) systems, which provide sequence-specific immunity by targeting foreign DNA (1–3). Other important strategies include abortive infection (Abi) systems (e.g., AbiD, AbiJ and Lamassu-Fam), which trigger programmed cell death to prevent phage spread (1, 4–7), and epigenetic interference systems (e.g., BREX and DISARM), which target and restrict phage replication (8). Recently, novel systems such as CBASS, Zorya, Pycsar, Druantia, and SspBCD-SspE (also known as DND system (9)) have been identified, each employing distinct signaling and enzymatic mechanisms to recognize and dismantle invaders (1, 4). For example, DND/SspBCD-SspE system protects bacteria by marking host DNA with phosphonothioate, allowing sspE to recognize and selectively cleave unmodified phage DNA (10). Additional systems like PD-T4-5, PD-T7-5, Shedu, and Gao_Qat further highlight the biochemical diversity of microbial defences (1–3, 7, 11–17). Collectively, these systems demonstrate the complexity and adaptability of bacterial immunity, reflecting an ongoing evolutionary arms race with phages.

Horizontal gene transfer (HGT) plays a fundamental role in bacterial evolution, enabling prokaryotes to rapidly acquire new genetic elements (18). Mobile genetic elements (MGEs) can impose costs on their host organisms and often spread selfishly within microbial populations. Nevertheless, they can also promote genome evolution and enhancement (19). However, their spread can be limited by bacterial defence mechanisms, such as those mentioned earlier (16). Interestingly, many MGEs, including bacteriophages and plasmids, often encode anti-defence proteins to overcome these barriers (3, 20). Although defence systems such as CRISPR-Cas protect bacteria from invading MGEs, they also incur fitness costs, potentially hindering beneficial genetic acquisitions, such as those conferring antibiotic resistance (21–23). While laboratory evidence supports the restrictive role of these systems, broader evolutionary studies present conflicting findings, reflecting the complex and context-dependent dynamics that vary across bacterial species and ecological niches (20, 24, 25).

*Acinetobacter* is a genus of gram-negative bacteria that cause opportunistic infections. Notably *Acinetobacter baumannii* is a major clinical threat due to its resistance to antibiotics, disinfectants, and environmental stressors, allowing it to persist on hospital surfaces and cause healthcare-associated infections (26). According to a systematic analysis, the multidrug resistant (MDR) *A. baumannii* is found to be associated with around 400,000 global deaths in 2019 (27). Its genomic plasticity, including resistance and defence islands, makes it a suitable model to study defence mechanisms and HGT (11, 28–31). Another species emerging globally, *Acinetobacter pittii*, carries a broad resistome within its accessory genome, raising concerns about its potential to follow the same high-risk, multidrug-resistant trajectory as *A. baumannii* (32). Other species such as *A. lwoffii*, *A. ursingii*, and *A. johnsonii* are increasingly recognized as nosocomial pathogens in ICUs, causing bloodstream, urinary tract, and wound infections, with multidrug resistance common in over two-thirds of isolates. Often underestimated, these non-baumannii *Acinetobacter* spp. show environmental persistence, biofilm formation, and resistance to disinfectants, enabling them to thrive in hospital settings and, therefore, demanding vigilant infection control (26, 32).

In this study, we analysed 132 genomes representing 18 *Acinetobacter* species. These genomes exhibited diverse geographical distributions and genomic characteristics, providing a robust foundation for investigating bacterial defence mechanisms. Given that most genomes were derived from clinical isolates, our dataset offered a unique perspective to explore the relationship between bacterial adaptation and selective pressures such as antibiotics and phage predation. We examined the distribution, diversity, and co-occurrence of defence systems and HGT markers, including antibiotic resistance genes (ARGs), heavy metal resistance genes (HMRGs), and integrative mobile elements (IMEs). We aimed to better understand how *Acinetobacter* spp. adapts to clinical environments under these selective pressures. We focused on the evolutionary trade-off between maintaining costly defence arsenals and acquiring beneficial genes through HGT. We compared our results against a collection of 90 *Acinetobacter baumannii* international clone 2 (IC2) genomes which is one of the most successful lineages of *A.Baumannii*. Notably, our findings show that the prevalence and composition of defence systems vary by species and environmental niche. We also identified both synergistic and antagonistic interactions between defence systems and HGT markers, highlighting the influence of ecological and evolutionary pressures on bacterial adaptive strategies. As phage therapy gains interest as an alternative to antibiotics, understanding the genetic basis of phage resistance in *Acinetobacter* is increasingly important. Our findings reveal a complex interplay among bacterial immunity, mobile genetic elements, viral threats, and resistance traits.

Based on these insights, we proposed a survival model for *Acinetobacter* in environments exposed to both antibiotics and phages. In this model, trade-offs between phage resistance, antibiotic tolerance, HGT burden, and the metabolic cost of maintaining defence systems are finely balanced, even after selective pressures decline. These insights may inform strategies to combat multidrug resistance and improve the success of phage-based treatments.

## Methods

### Genome collection

A dataset of n=122 diverse *Acinetobacter* spp. genome sequences were downloaded in FASTA format (accessed January,2025) from the NCBI *efetch* service using a custom bash script and a list of NCBI Accession Numbers (Supplementary Data S1, Sheet S2). This dataset primarily consisted a complete chromosome-level assemblies and represented broad taxonomic diversity across 18 *Acinetobacter* spp., allowing for comparative analyses across the genus. Additionally, we collected 100 *Acinetobacter baumannii* international clone 2 (IC2) genomes. The assemblies were further categorized based on genome assembly level: contig-level genomes (n=90) and complete genomes (n=10) (Supplementary Data S1, Sheets S3-S4). To preserve data integrity, the contig level assemblies (n=90) were analysed separately from complete chromosome assemblies (n=122+10=132) (Supplementary Figure 1, Supplementary Data S1, Sheet S18).

### Identification of defence and counter-defence system

All analyses were performed on Ubuntu v24.04.1 LTS. Bioinformatics software was managed with Miniconda v24.11.3, using the Bioconda and Conda-Forge channels enabled for package installation. Defence systems were predicted with DefenseFinder v1.3.0 (7) and PADLOC v2.0.0 (2). DefenseFinder, which relies on HMMER v3.4, and PADLOC, which integrates CRISPRCasFinder (33) for CRISPR array detection, were each installed in a dedicated Conda environment to ensure proper dependency management. Both tools were run in batch mode across all genomes using custom Bash scripts (available at: https://github.com/vikos77/Acinetobacter-defence-systems/tree/main/code/1_pipeline).

Output files were processed with command-line utilities (e.g., awk, sed) and consolidated into combined tab-separated summary files for each tool and dataset (Supplementary Data S1, Sheets S5-S11). Since both tools can detect same defence system multiple times within a single genome, additional hits were removed by deduplication in R using dplyr, to avoid overrepresentation of certain systems. Additional R packages (e.g., readr, tidyr, ggplot2, gridExtra) were used for data wrangling, statistical analysis, and visualisation (available at: https://github.com/vikos77/Acinetobacter-defence-systems/tree/main/code/2_analysis).

### Phylogenetic analysis of type-I BREX

Six type-I BREX proteins from various *Acinetobacter* plasmids were retrieved from the NCBI protein database (accessed June,2024). Functional annotations were then validated using HHPRED, and structural predictions were generated via the MPI Bioinformatics Toolkit under default parameters (https://toolkit.tuebingen.mpg.de/). The BrxR protein sequence was obtained from Luyten et al. (2022) (34). To identify homologs, PSI-BLAST searches (e-value cutoff 10^−6^) were performed, and the top 1000 hits were retrieved. Full-length homologous sequences were then clustered using MMseqs2 with default parameters, and multiple sequence alignment (MSA) was caried out using MUSCLE (35). A phylogenetic tree was then constructed using the maximum-likelihood method in MEGA12 (36) and visualized with iTOL v7 (37) (Supplementary Data S2).

### Identification of antibiotic resistance genes (ARGs)

Antimicrobial resistance (AMR) genes were identified using ResFinder v4.6.0 (38) with default command-line parameters (Supplementary Data S1, Sheet S12). Again, to ensure each genome was counted only once per resistance gene, deduplication step was performed in R using the dplyr package (available at https://github.com/vikos77/Acinetobacter-defence-systems/blob/main/code/2_analysis/4_resistance_gene_analysis.R).

### Identification of integrated mobile elements (IMEs)

IME protein sequences were retrieved from the ICEberg 2.0 database (39). Homologous IME proteins were identified through tblastn search (e-value < 1e-6; query coverage and sequence identity > 80%). Resulting hits were parsed to extract ICEberg element identifiers, protein functional annotations, and genomic coordinates. These data were then used for downstream correlation analysis with bacterial defence systems, using Fisher’s exact test with Benjamini-Hochberg correction for multiple testing (Supplementary Data S1, Sheet S14).

To eliminate redundancy from multiple protein accessions representing the same function, IME hits were grouped by protein function rather than individual ICEberg element–protein combinations. Uninformative annotations (e.g., hypothetical proteins, unknown functions, putative proteins) were filtered out to focus on characterised mobile genetic element functions. For correlation analysis with defence systems, the top protein functions were selected based on genome prevalence (i.e., the number of distinct genomes containing each function), rather than total hit counts, ensuring that each genome contributed equally to the analysis regardless of the number of mobile elements present.

### Identification of heavy metal resistance genes (HMRGs)

Heavy metal resistance genes (HMRGs) were retrieved from the BacMet database (40). To identify homologous HMRGs within the genomes analysed, a tblastn search was conducted. Stringent filtering criteria (e-value < 0.005; query coverage > 80%; sequence identity > 70%) were applied to ensure high-confidence matches (Supplementary Data S1, Sheet S13).

### Co-relation analysis between defence systems and HGT markers

To examine associations between defence/anti-defence systems and ARGs or IMEs or HMRGs, binary presence/absence matrices were generated. Fisher’s exact test was applied to each pair to calculate odds ratios and determine statistical significance. Results were visualized as bar charts of log_2_-transformed odds ratios and correlation heatmaps, with colour gradients indicating effect size and asterisks denoting significance (*p < 0.05, **p < 0.01, ***p < 0.001). To capture broader trends between defence system abundance and other genomic elements (ARG, HMRG, IME, and anti-defence) Spearman’s rank correlation was also applied, offering a non-parametric complement suited to count data and non-linear relationships (Supplementary Data S1, Sheets S16-S18).

## Results and Discussion

Analysis of 132 complete *Acinetobacter* spp. genomes presented broad geographical coverage across six continents, with highest representation from China (n=45), Mexico (n=13), and the USA (n=12) (Figure 1A). Over half of these genomes (55%) originated from human clinical samples, while environmental and unknown sources each constituted 17%, with faecal and animal-associated isolates contributing 8% and 4%, respectively (Figure 1B). The dataset comprised 43 *A. baumannii*, 27 *A. pittii*, and 62 isolates from 16 other species. In our opinion, the geographical and ecological diversity of the dataset, combined with the predominance of clinical isolates, provides a solid foundation for exploring the genomic basis of defence systems and pathogenicity in Acinetobacter, with our results offering key insights and supporting previous findings.

**Figure. 1:**
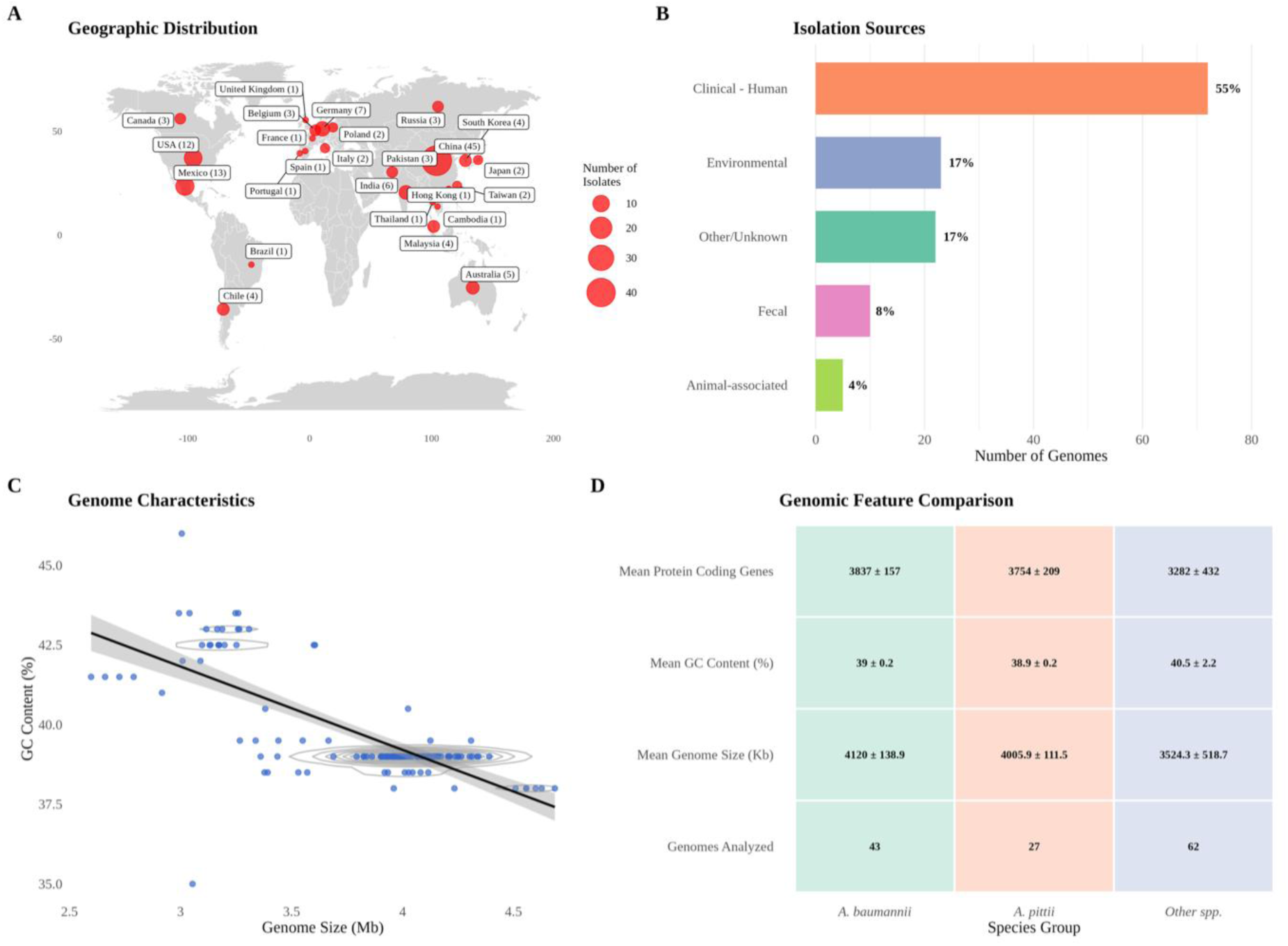
Epidemiological analysis of data. 132 genomes map making, source, GC vs genome size, protein coding gene. A: Global geographical distribution of *Acinetobacter* isolates, with colour intensity indicating relative abundance. All the countries are labelled with isolate numbers in parentheses. B: Distribution of isolation sources across the dataset categorized as Clinical - Human, Environmental, Others/Unknown, fecal, and Animal-associated sources. C: Relationship between genome size (Mb) and GC content (%). D: Comparative analysis of key genomic features across species groups, including genomes analysed, mean genome size, GC content, and protein-coding gene count for *A. baumannii* (n=43), *A. pittii* (n=27), and other *Acinetobacter* species (n=62).

Comparative genomic analysis showed significant inter-species variation in key genomic features such as genome size, GC content and protein-coding gene counts (Figure 1C-D). GC content varied significantly across species (F₂,₁₂₉ = 16.74, p < 0.001), but in this case, *A. baumannii* (39.0 ± 0.2%) and *A. pittii* (38.9 ± 0.2%) showed lower GC content than other species (40.5 ± 1.1%). Post-hoc analysis confirmed both clinical species harbouring significantly lower GC content from other Acinetobacter spp. (p < 0.001), with no difference between them (p = 1.0). One-way ANOVA indicated significant genome size differences among species (F₂,₁₂₉ = 37.59, p < 0.001), with *A. baumannii* (4120.0 ± 138.9 Kb) and *A. pittii* (4005.9 ± 111.5 Kb) possessing larger genomes than other Acinetobacter spp. (3524.3 ± 518.7 Kb). Post-hoc pairwise t-tests (Bonferroni corrected) confirmed these differences (p < 0.001), with no significant difference between *A. baumannii* and *A. pittii* (p = 0.63). Similarly, protein-coding gene counts also differed significantly (F₂,₁₂₆ = 41.44, p < 0.001), with *A. baumannii* (3837 ± 157) and *A. pittii* (3754 ± 209) encoding more genes than other species (3282 ± 432). Post-hoc tests confirmed significant differences between both clinical species and other Acinetobacter spp. (p < 0.001), but not between each other (p = 0.94).

The significantly larger genomes and higher gene counts in *A. baumannii* and *A. pittii*, compared to other species, could be associated with lineage-specific genome expansion driven by the acquisition of accessory elements, such as, genomic islands and mobile genetic elements (41). Also, lack of significant differences between these two species supports the idea of convergent adaptation among clinically important lineages (42). The inverse relationship between GC content and genome size further supports the role of HGT in genome evolution, as some previous findings linked lower GC content to acquisition of genomic islands and mobile genetic elements leading to genome expansion (41, 43, 44). These results highlight how genome architecture in Acinetobacter may be shaped by ecological context, influencing the distribution of defence system components.

### Niche-specific trade-Offs in defence system distribution

On average, the complete *Acinetobacter* genomes (n=132) in our dataset carried 4-6 defence systems (Figure 2A), consistent with the typical bacterial range of approximately 5–6 systems (14). Maintaining multilayered immunity entails trade-offs, including limited HGT and increased metabolic burden (14, 45). Evidence suggests that certain redundant or synergistic combinations, such as, methylation-Associated Defence System (MADS) working alongside CRISPR–Cas are more effective at preventing the spread of phage variants that can evade MADS alone (45–47). The widespread presence of multiple, mechanistically distinct systems in *Acinetobacter* likely represents an evolutionary balance, optimizing protection against phage attack while managing the associated costs. Strikingly, *A. baumannii* international clone 2 (IC2) (n=90) carried only 1–3 defence systems on average, much lower than other *A. baumannii* strains, which typically encoded around 5 (Figure 3A). IC2 has previously been characterized by an exceptionally high load of antibiotic resistance genes (ARGs), with an average of 17.1 per genome and resistance predicted against critical drugs such as carbapenem, sulbactam, and minocycline (31, 48). Found in over 60 countries (31), IC2’s epidemiological success may be partly linked to its reduced defence repertoire, which likely facilitates unrestricted HGT, enabling the rapid acquisition of resistance traits. This highlights the divergent evolutionary strategies within *Acinetobacter*, where some lineages favor defence-heavy genomes, while others, like IC2, appear to prioritize genetic plasticity for survival and global dissemination.

**Figure. 2:**
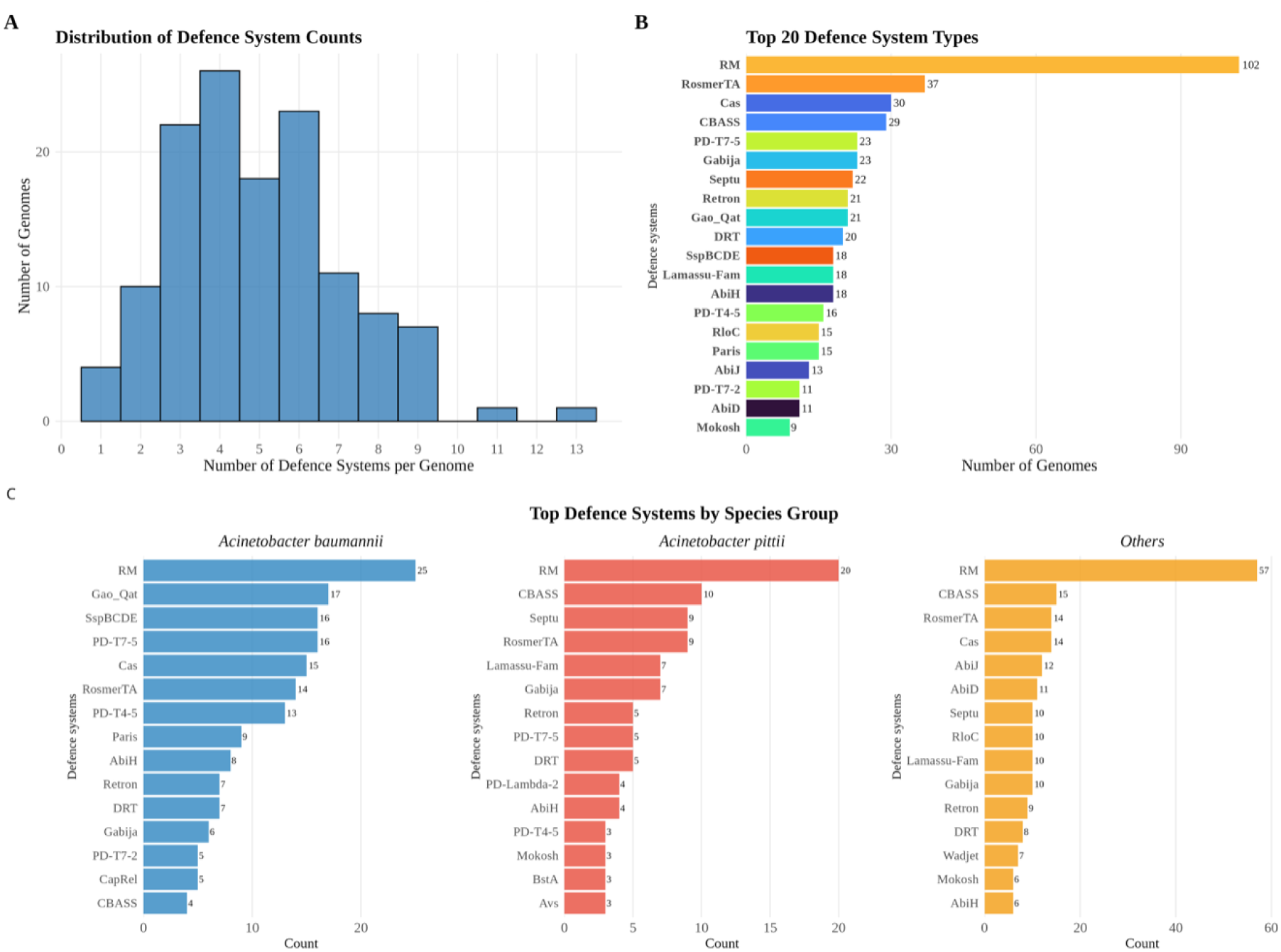
Genus-wide versus species-specific analysis of bacterial defence systems in *Acinetobacter spp*. **A** Distribution of defence system counts per genome across 132 complete *Acinetobacter* genomes. The histogram shows the frequency distribution of total defence systems detected per genome using DefenseFinder, with most genomes containing 4-6 defence systems. **B** Prevalence of the top 20 defence system types detected across the complete dataset. Restriction-Modification (RM) systems dominate the *Acinetobacter* defence landscape, present in 102 genomes (77.3%), followed by RosmerTA (37 genomes, 28.0%), and Cas systems (30 genomes, 22.7%). Numbers adjacent to bars indicate the absolute count of genomes containing each system type. **C** Species-specific distribution of defence systems across three taxonomic groups: *A. baumannii* (n=43), *A. pittii* (n=27), and other *Acinetobacter* species (n=62). While RM systems maintain high prevalence across all groups, distinct species-specific patterns emerge. *A. baumannii* shows enrichment of Gao_Qat (17 genomes) and SspBCDE (16 genomes), while *A. pittii* exhibits predominance of CBASS (10 genomes) and Septu (9 genomes) systems with notable absence of SspBCDE enrichment. Other *Acinetobacter* species display the highest diversity with prevalent CBASS (16 genomes) and RosmerTA (14 genomes) systems.

**Figure. 3:**
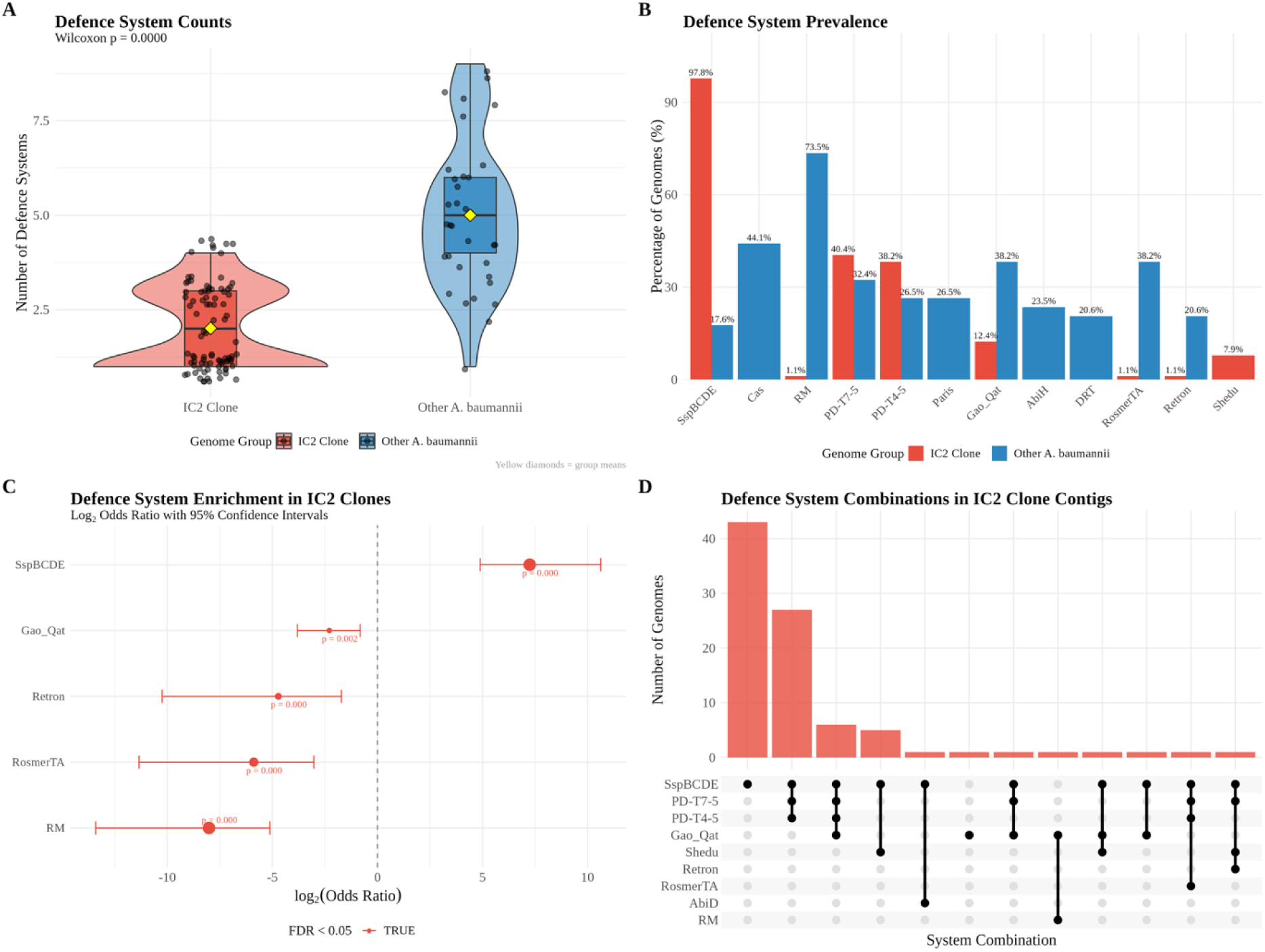
IC2 clone contigs analysis emphasizing on *A. baumannii*. **A** Comparative distribution of defence system counts between IC2 clone (n= 90) and other *A. baumannii* strains (n=43). Violin plots reveal that IC2 clones harbor significantly fewer defence systems per genome compared to other *A. baumannii* lineages (Wilcoxon rank-sum test, p = 0.000). IC2 clones show a narrow distribution centered around 1-3 systems, while other *A. baumannii* strains display greater diversity with 4-6 systems per genome. **B** Species-level comparison of defence system prevalence between IC2 clones (red bars) and other *A. baumannii* strains (blue bars). While SspBCDE systems show a major increase in prevalence between groups (97.8% vs 17.6%), IC2 clones exhibit notable depletion in most other defence systems, including RM (7.6% vs 73.5%), PD-T7-5 (24.2% vs 32.4%), and Gao_Qat (7.4% vs 38.2%). Percentages indicate the proportion of genomes within each group containing the respective system. **C** Statistical enrichment analysis of defence systems in IC2 clones relative to other *A. baumannii* strains. Forest plot displays log₂ odds ratios with 95% confidence intervals, revealing significant depletion (negative odds ratios) of multiple defence systems in IC2 clones, including Gao_Qat, Retron, RosmerTA, and RM systems (all FDR-corrected p < 0.001, indicated in red). **D** Defence system combination patterns in IC2 clone contig assemblies (n=90). The majority of IC2 clones contain SspBCDE as a single defence system (43 genomes), with other genomes harboring complex multi-system combinations. Black circles indicate presence of each system type, demonstrating the simplified defence architecture characteristic of this globally successful clinical lineage.

Our comparative analysis also revealed a striking difference in defence system distribution across lineages. Restriction–modification (RM) systems (14) were the most prevalent defence type overall present in 102 of 132 genomes (Figure 2B), nearly absent in IC2 (only 1.1%). Instead, the phosphorothioation system (known as DND/SspBCDE) (9–11, 49) was found in 97.8% of IC2 genomes, compared to 17.6% in others (Figure 3B-C). Although SspBCDE was the third most prevalent system in *A. baumannii* complete genomes (n=43), it was rare in *A. pittii* and other *Acinetobacter* species analyzed (only 2 out of 89 non-*A. baumannii* genomes; Figure 2C and Supplementary data 1), indicating lineage-specific selection. The enrichment of SspBCDE in *A. baumannii* may serve as a potential functional alternative to RM, offering selective immunity without restricting gene uptake and its rarity in other lineages implies strong niche-specific selection.

Several defence systems were mutually exclusive between IC2 and others. CRISPR–Cas, AbiH, and PARIS were entirely absent from IC2 but present in other lineages. Toxin–antitoxin systems such as RosmerTA and Retron were also rare in IC2 but more common in others (Figure 3B). In contrast, DNA-modification systems PD-T7-5 and PD-T4-5 were moderately present in both groups, suggesting conserved components in anti-phage defence.

Previous study reported that only ∼14% of *A. baumannii* strains possess CRISPR arrays, Cas genes, or both, with no significant link to antibiotic resistance genes (24). Our analysis using CRISPRCasFinder, PADLOC, and DefenceFinder revealed that complete CRISPR-Cas systems are indeed not very common in *Acinetobacter* Spp.: only 29 out of 132 genomes (22%) contained both CRISPR arrays and Cas genes, mostly of the type I-F subtype. The majority (63.6%) harboured only CRISPR arrays, indicating widespread non-functional or degenerate loci. Just one genome (0.8%) had *cas* genes without an array, underscoring the rarity of intact systems (Supplementary Data 1; Supplementary Figure 2A). Despite the apparent erosion of CRISPR-Cas systems, alternative defences were common. Among the 18 genomes (13.6%) lacking both arrays and *cas* genes, all carried other defence systems; primarily RM (77.8%), followed by RosmerTA (33.3%), AbiD (27.8%), AbiJ (22.2%), and Gabija (16.7%), highlighting a layered and diversified phage defence strategy (Supplementary Figure 2B). Strikingly, genomes with intact CRISPR-Cas systems (n = 29) encoded significantly more total defence systems (mean = 6.34 ± 2.41) than those with only CRISPR arrays (4.64 ± 1.67, p = 0.002) or lacking CRISPR-Cas entirely (4.28 ± 2.05, p = 0.013), based on Kruskal–Wallis analysis (p = 0.0015) (Supplementary Figure 2C). Genomes with only CRISPR arrays were statistically indistinguishable from those entirely lacking CRISPR–Cas loci (p = 1.0), underscoring the functional irrelevance of orphan arrays. Together, these findings suggest that while functional CRISPR–Cas loci enhance phage resistance, they remain susceptible to phage-encoded anti-CRISPR proteins (3). To confer immunity, bacteria appear to retain complementary systems (14). Conversely, the degradation of CRISPR–Cas machinery may facilitate horizontal gene transfer by lifting barriers to foreign DNA uptake, with other defence systems compensating for the lost adaptive function. The widespread occurrence of orphan CRISPR arrays likely reflects remnants of previously active systems, rather than ongoing roles in host defence (6, 9, 11, 14, 24, 25, 50).

Overall, these patterns reflect the complex interplay of evolutionary forces including horizontal gene transfer, recombination, and niche-specific selection that shape the diversification of defence arsenals in *Acinetobacter*. The high variability in gene content and organization within defence islands underscores the genus’s genomic plasticity, enabling rapid adaptation to phage pressure, antimicrobials and environmental stress (11). This flexibility may be a key factor in the clinical persistence and antimicrobial resistance frequently associated with IC2.

### Mutually exclusive defence systems shape the immune landscape

In line with our findings mentioned earlier, our co-occurrence analysis revealed a clear pattern of mutual exclusivity between restriction–modification (RM) systems and the SspBCDE system (Figure 4A-B). SspBCDE frequently co-occurs with other defence systems such as PD-T4-5, Gao_Qat, and PD-T7-5, but these combinations notably exclude RM systems. While both RM and SspBCDE act against foreign DNA, they employ distinct self vs non-self recognition strategies: methylation and phosphorothioation, respectively (6, 9, 14). This suggests that Acinetobacter species favour the assembly of complementary, rather than functionally redundant, defence systems to optimise resource use and broaden immune coverage (14).

**Figure. 4:**
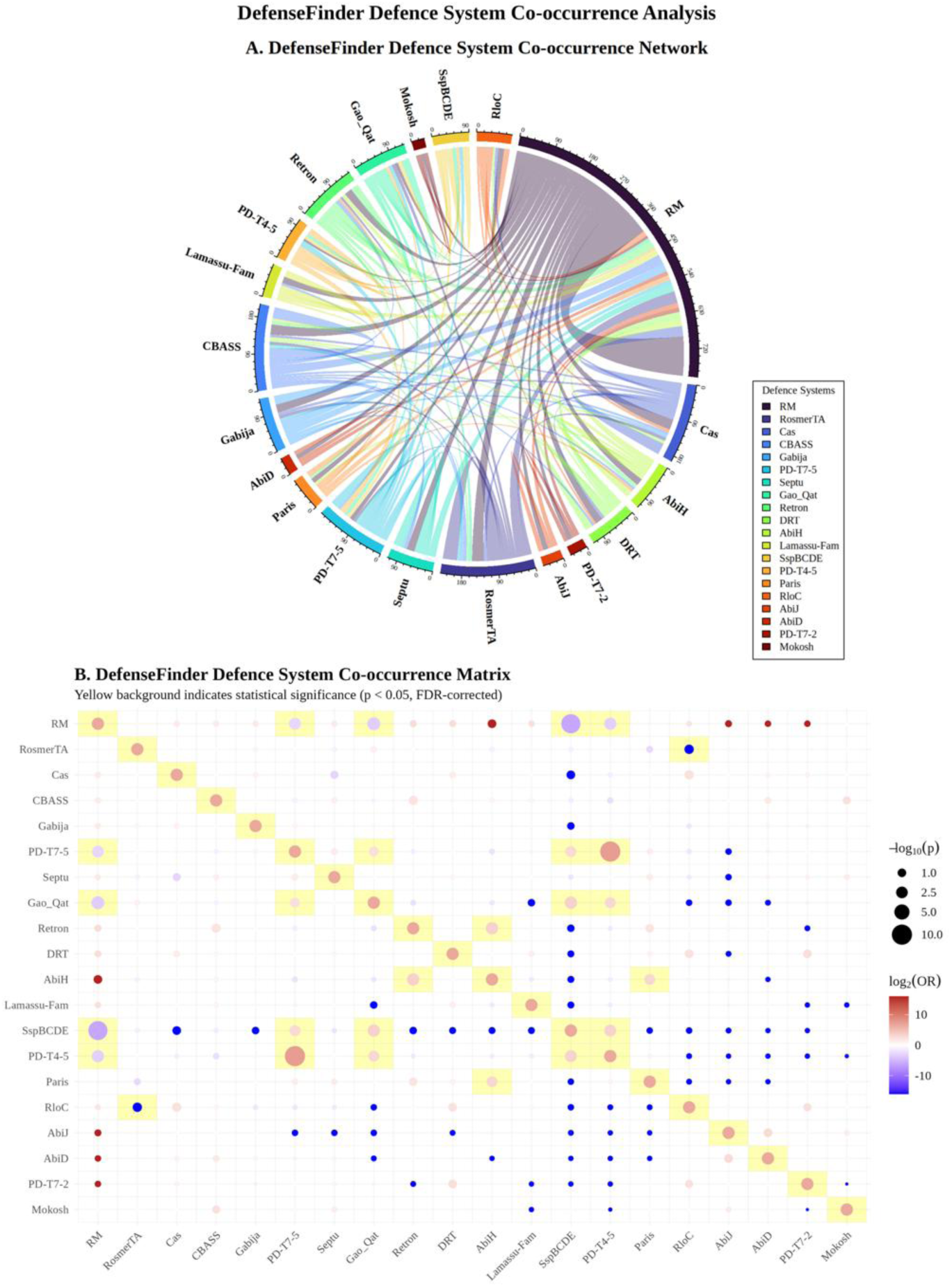
Defence system co-occurrence patterns. **A** Network visualization of defence system co-occurrence patterns across 132 complete *Acinetobacter* genomes. The circular chord diagram displays connections between defence systems, with line thickness and opacity representing the strength of association. RM systems (dark purple) demonstrate extensive connectivity with multiple other systems, while specialized systems like CBASS (blue) display broad connectivity while systems like Retron (green) and Gao_Qat (teal) show more selective association patterns. The network reveals both cooperative relationships (systems that frequently co-occur) and potential competitive exclusions between different defence strategies. **B** Statistical correlation matrix of pairwise defence system associations. Circle size represents the magnitude of association strength (log₂ odds ratio), while colors indicate the direction of association: red circles denote positive correlations (systems likely to co-occur), blue circles represent negative correlations (mutually exclusive systems), and yellow background highlights statistically significant relationships (p < 0.05, FDR-corrected). Strong positive correlations are observed between SspBCDE and other systems such as PD-T7-5, PD-T4-5, and Gao_Qat. Notable positive associations include Gao_Qat with PD-T7-5 systems, and CBASS with Retron systems. Conversely, strong negative associations are observed between RM systems and several specialized systems including PD-T7-5 and Gao_Qat.

A negative correlation was also observed between the presence of RloC tRNA-cleaving nuclease (51) and RosmerTA systems (15) (Figure 4B). The inverse correlation may reflect functional incompatibility, potentially due to overlapping targeting of nucleic acid substrates or conflicting regulatory pathways that prevent their stable co-maintenance within the same genome (52, 53). In contrast, significant positive correlations among Paris, AbiH, and retron defence systems (Figure 4B) suggest cooperative interactions, possibly layered or sequential immune responses, that afford Acinetobacter species greater flexibility in combating diverse phage threats (4, 7, 14, 45, 47). These findings warrant targeted experimental validation to elucidate the molecular basis of these interactions and to clarify their ecological and evolutionary significance.

In *A. baumannii* IC2 strains, the SspBCDE system emerges as a predominant standalone defence mechanism, present in approximately 97.8% of the analysed genomic contigs (Figure 4D). Its frequent integration into modular tripartite assemblies with PD-T4-5 and PD-T7-5, and associations with Shedu, Gao_Qat, and AbiD, highlight an adaptable immune architecture. This modularity likely reflects selective pressures favouring flexible and dynamic defence repertoires, enabling rapid response to environmental challenges such as phage infection and antibiotic stress.

### Type-I bacteriophage exclusion (BREX) system are common on plasmids but rarely found in chromosomes

To investigate the extra-chromosomal distribution of defence systems in *Acinetobacter* spp., we expanded our analysis beyond chromosomal sequences and included 283 plasmids associated with these 132 assembled genomes deposited in the NCBI database. Defence systems were detected only on ∼25% (69/283) plasmids. None encoded more than five systems, and over half (58%, 40/69) carried only a single system (Supplementary Figure 3A). In line with the chromosomal observation, restriction-modification (RM) systems predominated among 36% plasmids (25/69; Supplementary Figure 3B), underscoring their widespread conservation across genomic compartments. Interestingly, while our chromosomal analysis identified only one instance of Type I bacteriophage exclusion (BREX) system across 132 genomes (Supplementary data 1), plasmid exhibits notable enrichment with BREX system detected in 36% of the plasmids analyzed (25/69; Supplementary Figure 3B). Among these plasmids, 3 out of 25 harboured incomplete BREX gene clusters, lacking one or more of the seven canonical BREX components (Supplementary data 1). Typically, the BREX system includes six core genes (*pglX*, *pglZ*, *brxA–C*, *brxL*), often accompanied by another regulatory gene *brxR*, encoding a WYL domain protein. It confers protection against phages through DNA methylation without direct cleavage, thus preserving the integrity of the host genome (13, 14, 17).

Sequence alignments indicated low similarity between chromosomal and plasmid-encoded BREX proteins, with plasmid variants notably shorter. A comprehensive phylogenetic analysis incorporating all BREX proteins available in the NCBI database confirmed their evolutionary relatedness, except for BrxB. Interestingly, plasmid-encoded BrxB variants clustered with chromosomal BREX systems from other organisms rather than those from Acinetobacter, suggesting divergent evolution or functional specialisation (Supplementary Figure 4).

The preferential plasmid localisation of BREX suggests an adaptive strategy whereby bacteria enhance phage resistance by linking immunity with mobile genetic elements. This may promote co-transfer of phage defence and antimicrobial resistance, especially in clinical settings where both antibiotics and phages apply selective pressure. These findings align with recent reports of plasmid-encoded BREX systems co-occurring with resistance genes in zoonotic pathogens such as *Escherichia fergusonii* (13, 34), highlighting the clinical significance of this mobile defence loci.

Additional plasmid-borne systems identified include Mokosh, cyclic oligonucleotide-based anti-phage signalling systems (CBASS), and PD-T4-5 (Supplementary Figure 3B-C), further supporting the view that plasmids serve as reservoirs for auxiliary defence mechanisms (54). Remarkably, BREX and RM systems were mutually exclusive on the same plasmid, mirroring the antagonistic SspBCDE–RM pattern observed chromosomally (Supplementary Figure 3D).

### Defence systems-HGT markers correlations suggest complex adaptive strategies

The correlation matrix revealed complex relationships between defence systems and HGT markers, challenging the notion of a simple antagonistic relationship (Figure 5A). In general, defence systems showed weak negative correlations with antibiotic resistance genes (ARGs) (r = −0.11), integrative mobile element (IME) functions (r = −0.01), and heavy metal resistance genes (HMRGs) (r = −0.13). These findings are consistent with the canonical role of defence systems in limiting foreign DNA integration and thereby constraining HGT (3, 4, 6, 12, 14, 24, 50, 52, 55, 56). Conversely, strong positive correlations were observed between mobile genetic elements and resistant traits. For instance, ARGs and IME-related proteins were positively associated (r = 0.70, p < 0.001), strongly supporting the role of mobile elements as major vectors for resistance gene dissemination [46, 47].

**Figure. 5:**
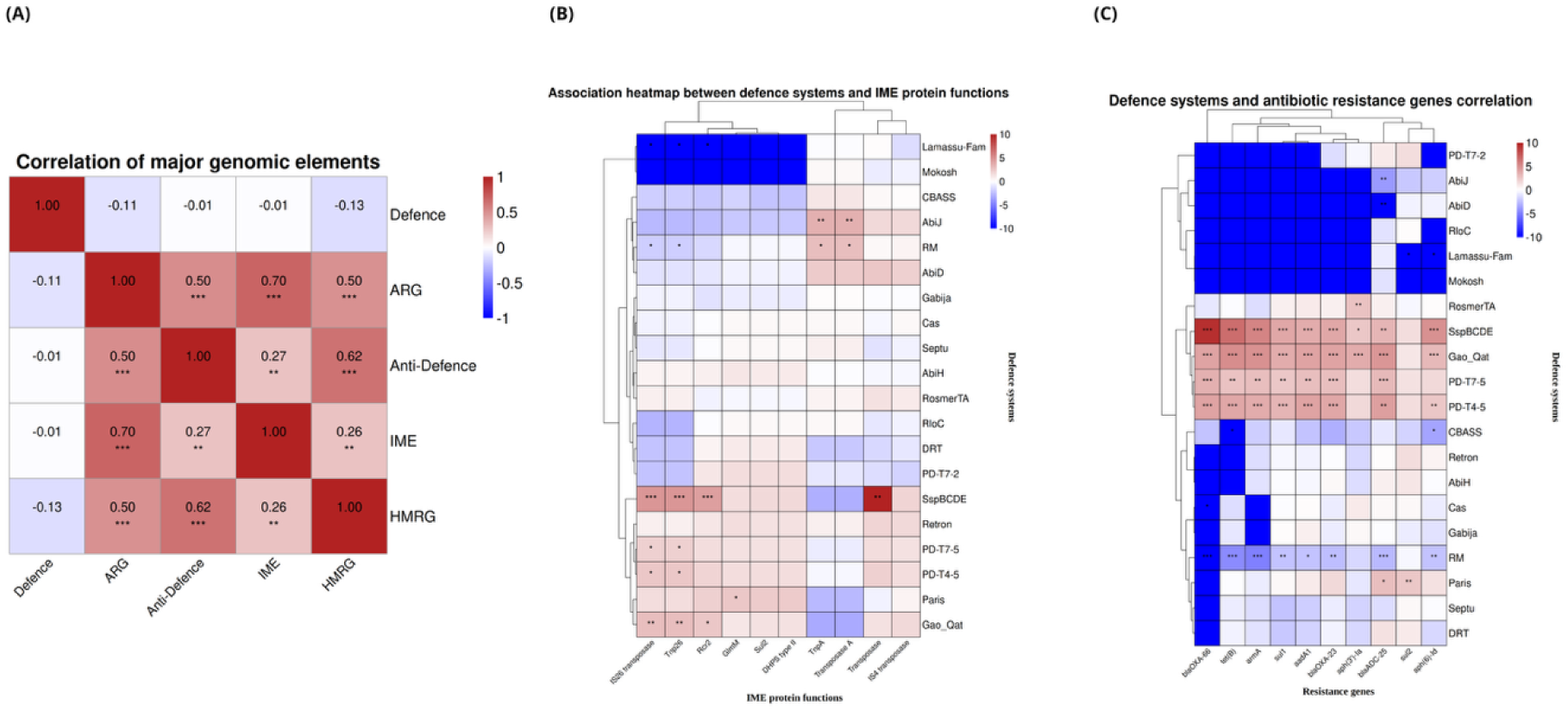
Complex interplay between defence systems, mobile genetic elements, and antibiotic resistance. **(A)** Correlation matrix between major genomic element categories across 132 *Acinetobacter* genomes. The 5×5 correlation matrix displays relationships between defence systems, antibiotic resistance genes (ARGs), anti-defence systems, integrative mobile elements (IMEs), and heavy metal resistance genes (HMRGs). Color scale represents correlation strength: dark red indicates strong positive correlations, white represents no correlation, and blue denotes negative correlations. Strong positive correlations supporting co-selection theory include ARGs and HMRGs (r = 0.50, p < 0.001), ARGs and IMEs (r = 0.70, p < 0.001), and HMRGs and anti-defence systems (r = 0.62, p < 0.001). HMRGs also show significant positive correlation with IMEs (r = 0.26, p < 0.01), indicating potential co-mobilization of metal and antibiotic resistance determinants. Defence systems show weak negative correlations with most mobile elements, suggesting restriction of horizontal gene transfer. **(B)** Association heatmap between defence systems and IME protein functions using hierarchical clustering. Color intensity reflects association strength: red indicates positive associations (systems likely to co-occur with specific IME functions), blue represents negative associations (mutually exclusive relationships), and white indicates no significant association. Notable positive associations (red clusters) include PD-T7-5, Gao_Qat, and SspBCDE systems with multiple IME protein functions, while Lamassu-family and RM systems show predominantly negative associations (blue regions), consistent with their restrictive role in horizontal gene transfer. **(C)** Correlation patterns between defence systems and antibiotic resistance genes with bidirectional hierarchical clustering. Red coloring indicates positive correlations (co-occurrence), blue represents negative correlations (mutual exclusion), and intensity reflects association strength. Gao_Qat, SspBCDE, and PD-T7-5 systems demonstrate strong positive correlations (intense red) with multiple resistance genes including β-lactamases (blaOXA variants), tetracycline resistance (tetB), and aminoglycoside resistance genes. Conversely, RM systems show consistent negative correlations (blue regions) with most ARGs, reinforcing their role as barriers to resistance gene acquisition. Statistical associations were calculated using Spearman correlation (Panel A) and Fisher’s exact tests with Benjamini-Hochberg correction for multiple testing (Panels B and C). Significance levels: * p < 0.05, ** p < 0.01, *** p < 0.001.

The function-based analysis of mobile element proteins revealed specific mechanisms underlying these resistance gene associations (Figure 5B). Transposases, particularly those associated with IS26 and Tnp26, positively correlated with synergistic defence systems (e.g., SspBCDE, PD-T&-5, PD-T4-5, Gao_Qat) suggesting possible co-mobilization of immunity and resistant traits. Rolling circle replicases (Rcr2), which are essential for plasmid replication and maintenance (57, 58), were also positively associated with SspBCDE and Gao_Qat, suggesting that certain immune mechanisms may stabilise resistance-carrying mobile elements (59).

Our analysis of HMRGs across *Acinetobacter spp.* provided further insights into resistance evolution. *A. baumannii* exhibited the greatest diversity of HMRGs and ARGs, supporting the co-selection theory where metal exposure indirectly promotes antibiotic resistance (60, 61). We observed significant positive correlation between ARGs and HMRGs (r = 0.50, p < 0.001) (Figure 5A) which supports this hypothesis. Additional positive associations between HMRGs and both anti-defence systems (r = 0.62, p < 0.001) and IMEs (r = 0.26, p < 0.01) indicate potential co-transfer of metal resistance, anti-defence, and antibiotic resistance determinants. This co-occurrence intensifies concerns over multidrug resistance, particularly in environments with high metal contamination. Importantly, clinical isolates exhibited significantly higher resistance gene counts than environmental strains (Supplementary Figure 5B), reflecting divergent selective pressures and niche-specific adaptation strategies.

Analysis of individual defence systems revealed marked contrasts in their associations with both ARGs and HMRGs, highlighting diverse immune strategies within *Acinetobacter* (Figure 5C and Supplement Figure 5C). Systems such as SspBCDE and Gao_Qat exhibited significant positive correlations (p < 0.001) with a broad range of β-lactamase and other resistance genes, including *blaOXA-66*, *tetB*, *armA*, and *sul1*. Notably, Gao_Qat was also positively associated with several HMRGs, including *adeG*, *abeS*, and *nreB* (Supplementary Figure 4C), indicating that these systems may have evolved permissive strategies that allow the selective acquisition of advantageous resistance traits while retaining anti-phage defence functions (55). Interestingly, *nreB*, which encodes a Ni²⁺/H⁺ antiporter involved in nickel resistance and is reported to be transferred to different bacterial species through HGT (60, 62–64), fits in line with our hypothesis. Similarly, the PD-T7-5 and PD-T4-5 systems showed strong positive associations with HMRGs like *adeG*, *abeS*, and *abuO* (Supplementary Figure 5C), alongside their established correlations with antibiotic resistance genes, indicating they also facilitate or at least do not restrict broad-spectrum resistance acquisition. However, restriction-modification (RM) systems consistently exhibited strong negative associations with both antibiotic and heavy metal resistance gene panels, with significant negative correlations (p < 0.001) observed across most resistance determinants tested. The AbiD and AbiJ systems similarly displayed negative associations with multiple ARGs and HMRGs, reinforcing their restrictive role against foreign DNA integration.

Overall, our findings point to a nuanced relationship between bacterial defence systems and the mobilome. Rather than a purely antagonistic dynamic, this relationship appears to be shaped by a complex interplay of selective forces. While canonical systems such as RM remain barriers to HGT, other systems such as Gao_Qat may facilitate the selective acquisition of mobile genetic elements under antibiotic and phage pressure. This highlights the need for further exploration into the mechanisms and selective pressures that enable such mobility and potential fitness benefits under environmental stress.

## Conclusion

In this study, we comprehensively analysed defence system architectures across *Acinetobacter* species, shedding light on their genomic diversity and ecological adaptations. Our findings highlight the distinctive defence landscape of the clinically dominant *A. baumannii* IC2 clones, which exhibit a notable depletion of classical immune systems alongside a marked enrichment of the phosphorothioation-based SspBCDE system. Although consistent with previous large-scale study in *A. baumannii* (11), our focus on IC2 strains offers new insights into clone-specific evolutionary strategies.

We observed complex patterns of mutually exclusive and co-occurring defence systems, indicating a finely balanced trade-off between phage resistance, horizontal gene transfer, and antimicrobial resistance. Additionally, frequent plasmid-borne BREX systems and partially compromised chromosomal CRISPR-Cas loci suggest a strategic adaptation for gene acquisition while retaining basic phage immunity—a beneficial strategy under intense antibiotic and phage pressure in clinical settings (Figure 6). This adaptability likely aids bacterial survival and contributes to the persistence of multidrug-resistant strains in clinical environments.

**Figure. 6:**
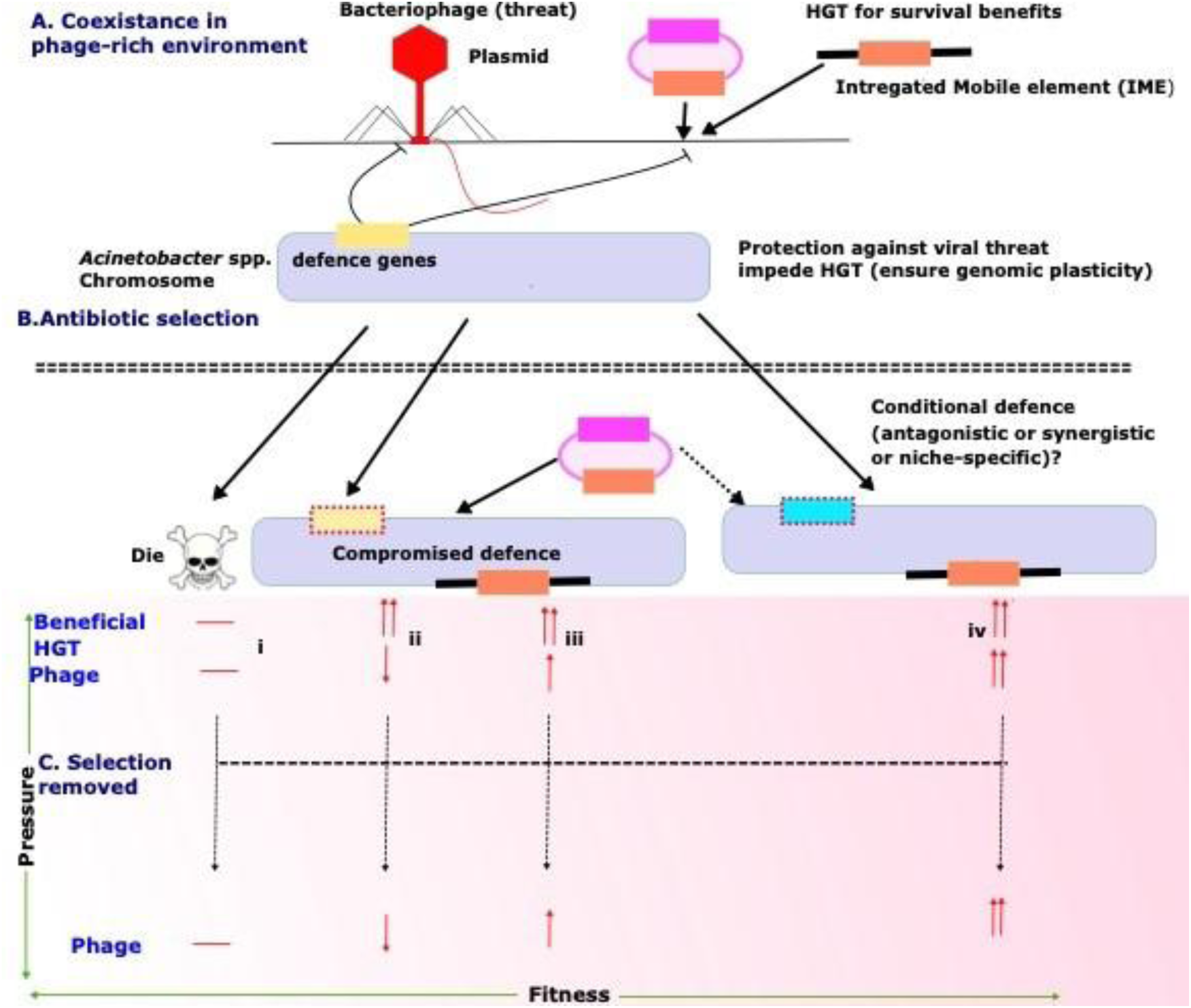
Proposed survival model of *Acinetobacter* spp. in a dual pressure environment composed of antibiotics and phages. **A** Schematic representation of an *Acinetobacter* chromosome carrying multiple defence loci (yellow). These systems can restrict phage infection, but at the same time may impede horizontal gene transfer (HGT) and thus limit acquisition of accessory DNA such as antibiotic-resistance genes (ARGs; orange) located on plasmids or other mobile genetic elements, preserving genomic plasticity. **B** Four potential fates of a bacterial cell exposed simultaneously to an antibiotic and a phage, with fitness changes denoted by upward (↑) or downward (↓) arrows: (i) Death (Die) – loss of viability due to antibiotic sensitivity. (ii) Compromised defence – mutation or deletion of defence genes (yellow with dashed border) improves fitness under antibiotic selection by facilitating ARG uptake but reduces fitness during phage predation. (iii) Plasmid acquisition – uptake of a plasmid that co-encodes an ARG and a phage-defence module (purple) restores fitness under both antibiotic and phage pressures, offsetting the metabolic cost of defence maintenance. (iv)Conditional defence expression (cyan with dashed border)– environment-dependent, mutually exclusive, synergistic, or niche-specific activation of defence systems (cyan with dashed borders) optimizes the trade-off between phage resistance, antibiotic tolerance, HGT burden, and the metabolic cost of maintaining defence components, even after selective pressures are removed.

These findings offer a foundation for the rationally designing targeted phage therapies against multidrug-resistant Acinetobacter. Although our dataset includes over 200 genomes (including 90 IC2 contigs) from 18 species, future studies with larger and more diverse genome collections are necessary to validate and expand upon our observations. Moreover, experimental validation of the synergistic and antagonistic interactions among defence systems and their regulation under various environmental and clinical conditions remains a critical next step. Ultimately, the integration of genomic surveillance with functional studies and synthetic phage engineering holds great promise for developing next-generation antibacterial strategies. Such efforts will be crucial in delivering precision-targeted therapies that remain effective amidst the ongoing evolution of antimicrobial resistance.

## Conflicts of interest

Dr Bruno Silvester Lopes is a member of the Editorial board for Journal of Medical Microbiology.

## Funding information

This work received no specific grant from any funding agency.

## Author contributions

Conceptualization: V.M. and S.S.; Formal analysis: V.M., S.S., P.D. and P.R.; Supervision: S.S.; Validation and visualization: V.M., P.R., and S.S.; Writing – original draft: V.M., S.S., B.S.L., and P.R.; Writing – review and editing: all authors.

V.M. and P.R. contributed equally to this work.

## Notes

https://github.com/vikos77/Acinetobacter-defence-systems

### Competing Interest Statement

The authors have declared no competing interest.

