## Supplement Figures for "Adaptive trade-offs between niche-driven defence system selection and horizontal gene transfer suggests clinical success in *Acinetobacter* spp"

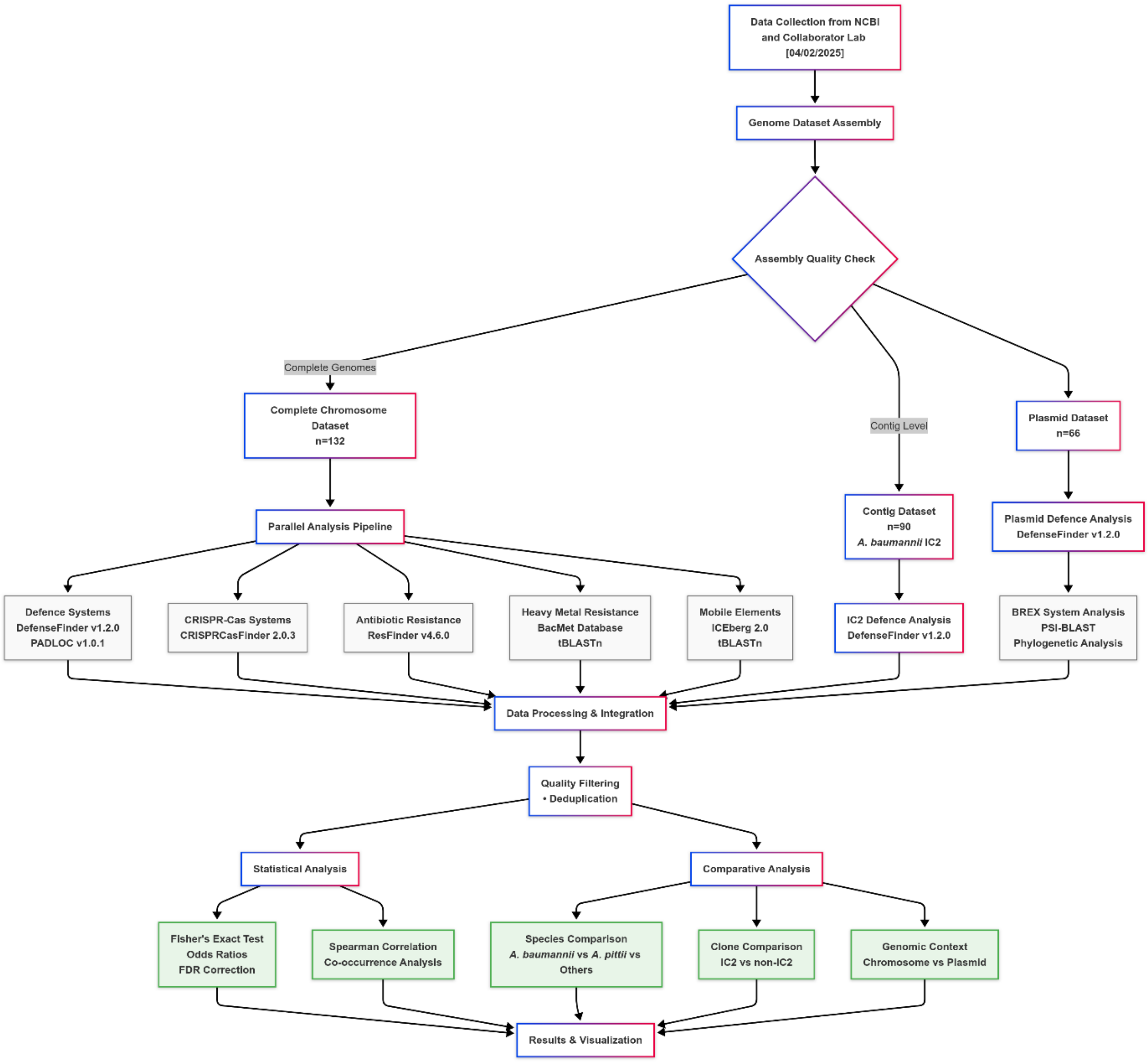


**Supplement Figure 1. Integrated workflow for defence system analysis in *Acinetobacter* spp.**

We analyzed 132 complete *Acinetobacter* genomes plus 90 *A. baumannii* IC2 contigs and 66 plasmids using parallel bioinformatics pipelines. Defence systems were identified with DefenseFinder/PADLOC, while resistance genes and mobile elements were analysed using ResFinder, BacMet, and ICEberg databases. After quality filtering and binary matrix generation, we applied Fisher's exact tests, Spearman correlations, and comparative analyses to reveal species-specific patterns, IC2-specific adaptations, and complex relationships between bacterial immunity and horizontal gene transfer. This comprehensive approach uncovered how clinical *Acinetobacter* strains balance phage resistance with genetic plasticity for survival in hospital environments.

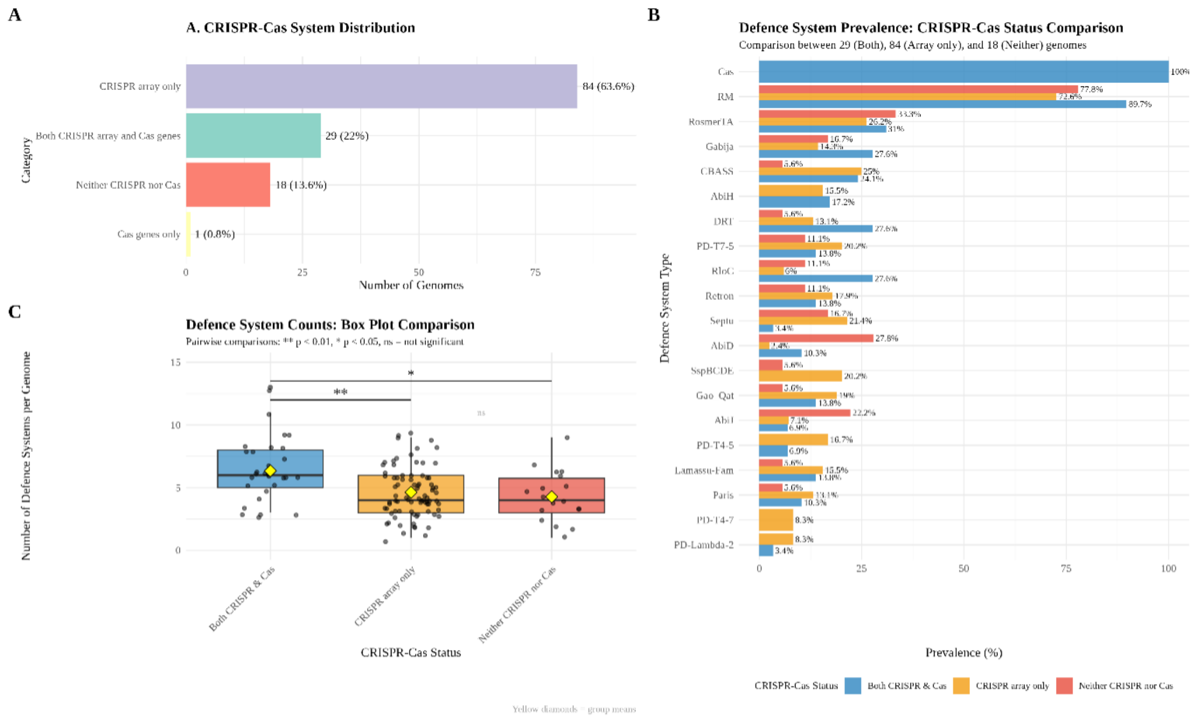


**Supplement Figure 2**: **Analysis of CRISPR-Cas distribution and its relationship with defence system prevalence across 132 *Acinetobacter* genomes.** (A) Distribution of CRISPR-Cas system categories showing predominance of non-functional arrays (63.6%) over complete systems (22%). (B) Defence system prevalence comparison across three CRISPR-Cas categories: functional systems (n=29), arrays only (n=84), and absent systems (n=18). Values show percentage of genomes containing each defence type. (C) Statistical comparison of defence system counts per genome across CRISPR-Cas categories. Box plots show median, quartiles, and individual data points; yellow diamonds indicate group means. Kruskal-Wallis test p = 0.0015; pairwise comparisons: ** p < 0.01, * p < 0.05, ns = not significant.

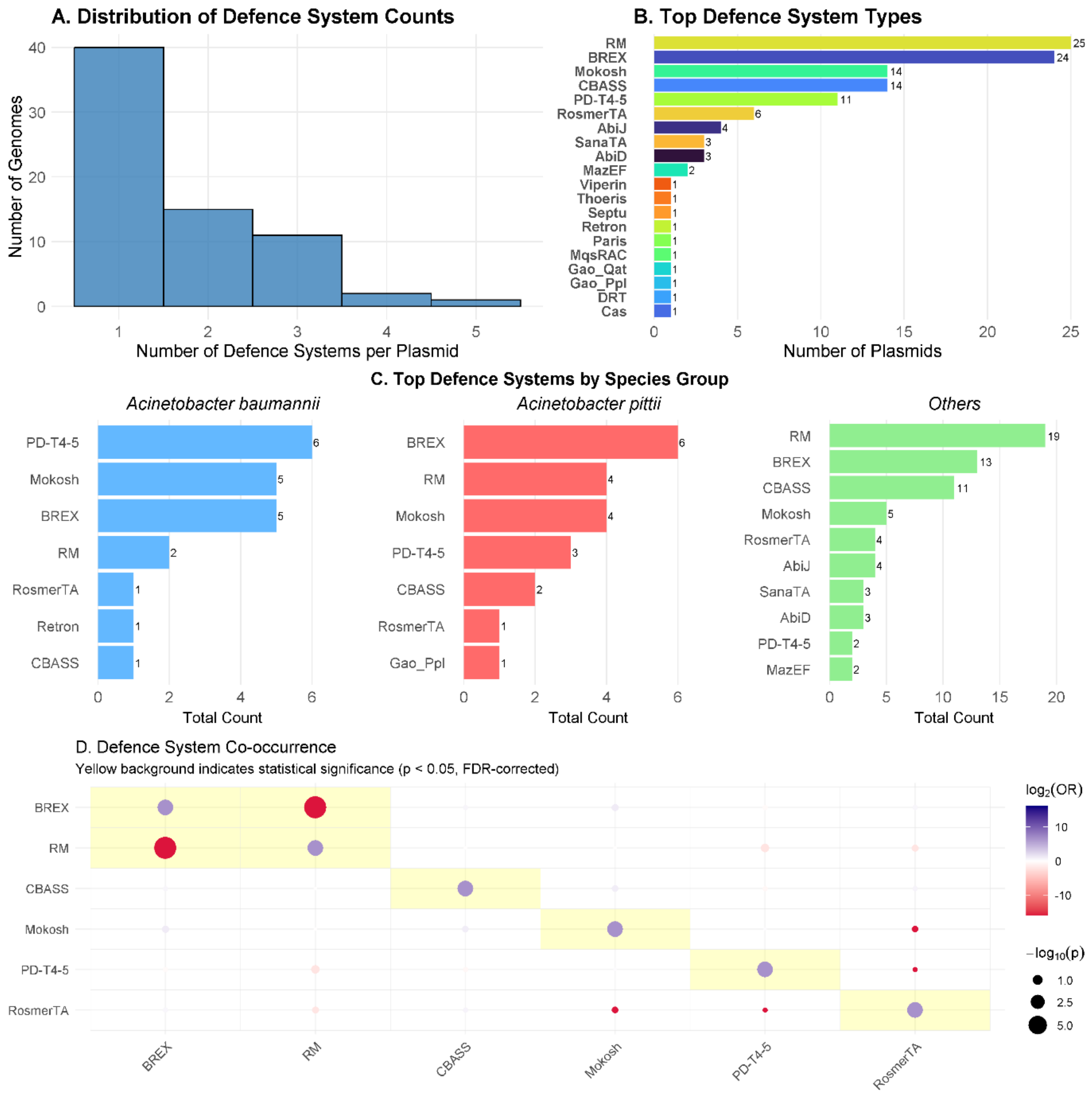


**Supplement Figure 3**: **Plasmid-borne defence systems in *Acinetobacter spp*. (A)** Distribution of defence system counts per plasmid across *Acinetobacter* species. The majority of plasmids carry single defence systems (35 plasmids), with decreasing frequency of plasmids harboring multiple systems. **(B)** Prevalence of defence system types detected on plasmids. Restriction-Modification (RM) systems dominate plasmid-borne defence mechanisms (26 plasmids), closely followed by BREX systems (25 plasmids). Secondary systems include Mokosh and CBASS (14 plasmids each), with numerous specialized systems present at lower frequencies. Numbers adjacent to bars indicate the absolute count of plasmids containing each system type. **(C)** Species-specific distribution of plasmid-borne defence systems across *A. baumannii* (n=7, blue bars), *A. pittii* (n=10, red bars), and other *Acinetobacter* species (n=49, green bars). *A. baumannii* plasmids show co-dominance of Mokosh and BREX systems (5 plasmids each), while *A. pittii* plasmids are enriched for BREX systems (6 plasmids). Other *Acinetobacter* species demonstrate strong preference for

RM systems on plasmids (20 plasmids). **(D)** Co-occurrence patterns of defence systems on plasmids. Circle size represents statistical significance level ($-\log_{10}(p)$), with larger circles indicating stronger significance. Blue circles denote negative associations (systems likely to co-occur on the same plasmid), red circles represent positive associations (mutually exclusive systems), and yellow background highlights statistically significant relationships ($p < 0.05$, FDR-corrected). Notable positive associations include BREX and RM, while some systems show competitive exclusion patterns.

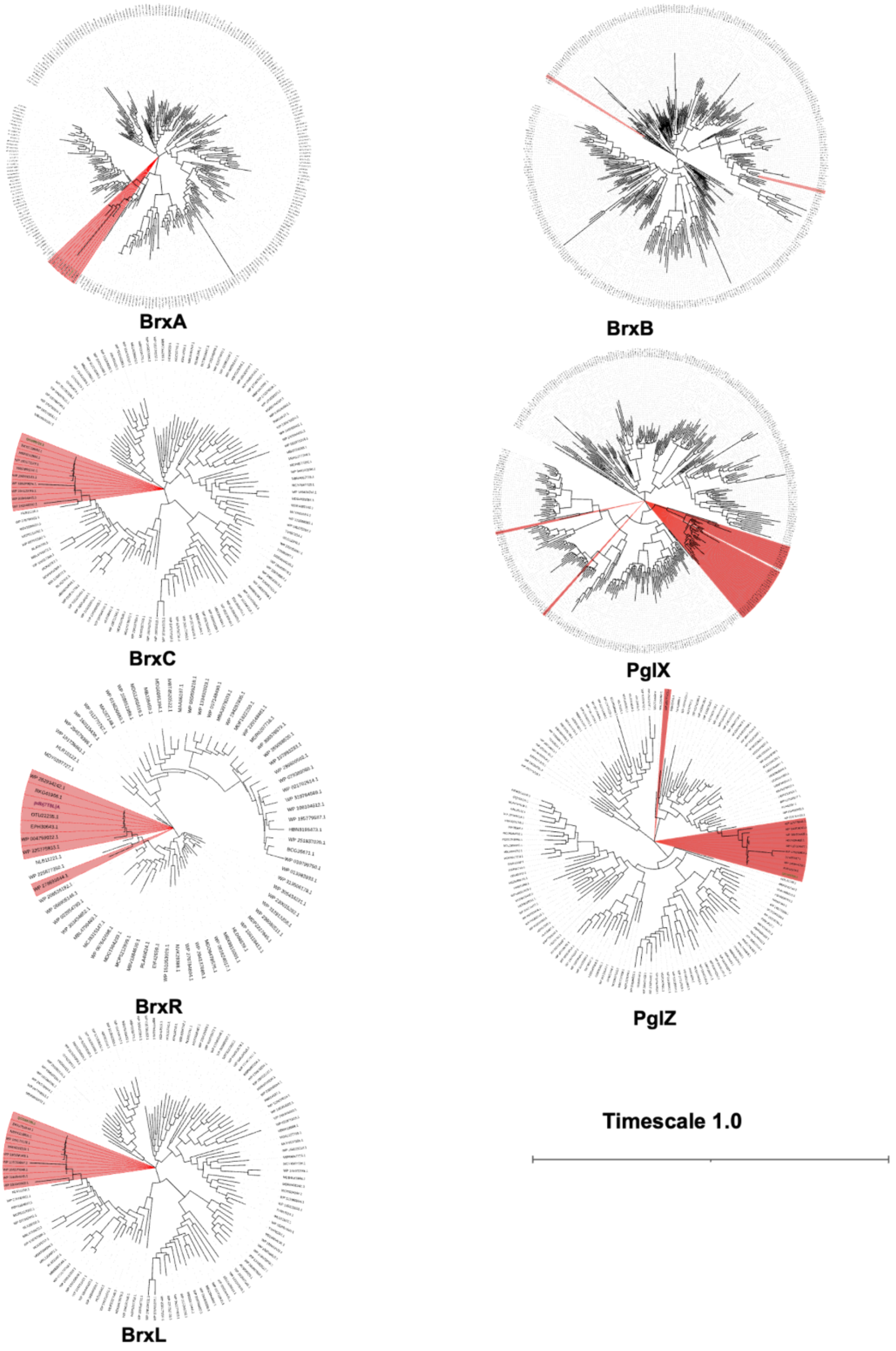

**Supplementary Figure 4**: **Phylogenetic distribution of Type-I BREX system proteins across bacterial species.** The phylogenetic tree depicts the evolutionary relationships of Type-I BREX system proteins BrxA, BrxB, BrxC, BrxR, BrxL, PglX, and PglZ among diverse bacterial taxa. Protein sequences originating from *Acinetobacter* spp. are highlighted in red, emphasizing their distinct lineage. Additionally, proteins encoded on plasmids are indicated with green bold text to distinguish plasmid-borne elements from chromosomal counterparts.

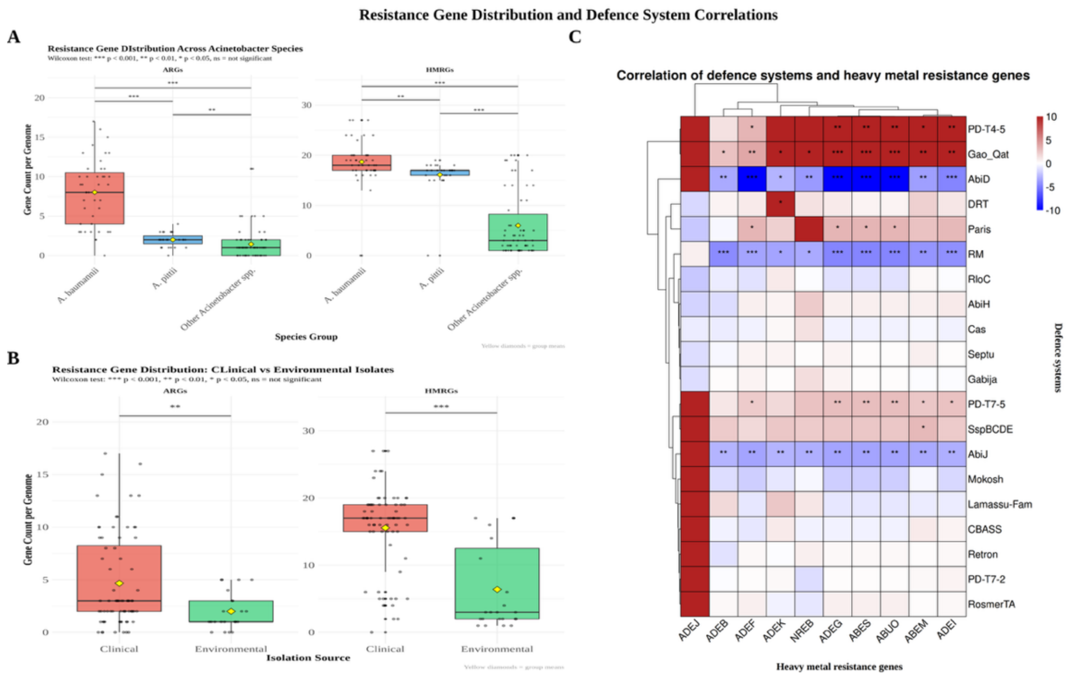


**Supplement figure 5**: **Resistance gene distribution and defence system correlations reveal species-specific and niche-adaptive patterns in *Acinetobacter* spp**. **(A)** Box plots comparing ARG and HMRG counts per genome across three taxonomic groups: *A. baumannii* (n=43), *A. pittii* (n=27), and other *Acinetobacter* species (n=62). *A. baumannii* demonstrates significantly higher loads of both resistance gene types compared to other species groups, supporting its clinical dominance and genomic plasticity. Yellow diamonds indicate group means, and individual data points are shown as overlaid dots. Wilcoxon rank-sum tests were performed for pairwise comparisons between groups. **(B)** Comparative analysis of ARG and HMRG prevalence between clinical (human-associated) and environmental isolates across the dataset. Clinical isolates show distinct resistance profiles compared to environmental strains, reflecting different selective pressures in hospital versus natural environments. Yellow diamonds indicate group means, and individual data points are shown as overlaid dots. Wilcoxon rank-sum tests were performed for pairwise comparisons between groups. **(C)** Heatmap displaying statistical associations between bacterial defense systems (y-axis) and individual HMRGs (x-axis) using bidirectional hierarchical clustering. Color intensity reflects $log_2$ odds ratio values: red indicates positive associations (defense systems likely to co-occur with specific HMRGs), blue represents negative associations (mutually exclusive relationships), and white shows no significant association. Notable positive associations include PD-T4-5, Gao_Qat, and PD-T7-5 systems with multiple

HMRGs, while RM systems show predominantly negative correlations with most HMRGs, suggesting restriction of horizontal gene transfer. Statistical associations were calculated using Fisher's exact tests with Benjamini-Hochberg correction for multiple testing across 132 complete Acinetobacter genomes. Significance levels are indicated by asterisks: * $p < 0.05$, ** $p < 0.01$, *** $p < 0.001$.
